# A single microfluidic device for multi-omics analysis sample preparation

**DOI:** 10.1101/2024.11.17.624005

**Authors:** Ranjith Kumar Ravi Kumar, Iman Haddad, Massamba Mbacke Ndiaye, Martial Marbouty, Joelle Vinh, Yann Verdier

**Affiliations:** Spectrométrie de Masse Biologique et Protéomique SMBP, ESPCI Paris, PSL University, LPC CNRS UMR 8249, 10 rue Vauquelin F-75005 Paris, France; Institut Pasteur, Université Paris Cité, Spacial Regulation of Genome Group, CNRS 3525 – 25-28 Rue du Dr Roux, F-75015 Paris, France

## Abstract

Combining different “omics” approaches, such as genomics and proteomics, is necessary to generate a detailed and complete insight into microbiome comprehension. Proper sample collection and processing and accurate analytical methods are crucial in generating reliable data. We previously developed the ChipFilter device for proteomic analysis of microbial samples. We have shown that this device coupled to LC-MS/MS can successfully be used to identify microbial proteins. In the present work, we have developed our workflow to analyze concomitantly proteins and nucleic acids from the same sample. We performed lysis and proteolysis in the device using cultures of *E. coli, B. subtilis*, and *S. cerevisiae*. After peptide recovery for LC-MS/MS analysis, DNA from the same samples was recovered and successfully amplified by PCR for the 3 species. This workflow was further extended to a complex microbial mixture of known compositions. Protein analysis was carried out, enabling the identification of more than 5000 proteins. The recovered DNA was sequenced, performing comparable to DNA extracted with a commercial kit without proteolysis. Our results show that the ChipFilter device is suited to prepare samples for parallel proteomic and genomic analyses, which is particularly relevant in the case of low-abundant samples and drastically reduces sampling bias.

## 1. Introduction

Microbiomes comprise many bacteria, viruses, fungi, and protozoa that interact in networks that vary spatially and with host states (1). The gut microbiome plays a role in various functions essential for health, including digestion, immune regulation, maintenance of the intestinal mucosal barrier, and protection from pathogens. Due to the importance of the microbiome in human physiopathology, this one is increasingly studied. The analysis of microbiomes should describe a complex web of interactions between microorganisms and their host. These interactions are difficult to characterize, underlining the crucial importance of multi-omics analyses (2).

Multi-omics analyses include various data layers that provide complementary information and must be approached with their considerations. These approaches include genomics, transcriptomics, proteomics, epigenomics, radiomics, metabolomics, lipidomic, diet, or clinical outcomes. Among these approaches, metagenomic is a method of choice to identify and quantify the microbiome species. On the other hand, metaproteomics enables functional activity information to be gained from the samples. Combining microbiome composition profiling with functional-omic platforms is attractive because it allows for a holistic overview of the functional state of the microbiome (3).

Sample preparation is a key point and constitutes a challenge in describing field samples using multiomics approaches, particularly for standardization (4). Traditionally, multiomics studies that combine genomics, transcriptomics, and proteomics analyses rely on separate sample preparation workflows for analysis5. Such split workflows can induce sample bias and tend to be labor-intensive and costly, relying on several separately purchased sample preparation kits or homemade protocols. For example, commercially available kits that allow the collection of DNA, RNA, and protein from a single starting sample include the TriplePrep kit (Cytiva) and the AllPrep DNA/RNA/Protein kit (Qiagen). This kit relies on separate columns to purify nucleic acids and yields intact proteins that must be precipitated before downstream processing for MS analysis (5).

In recent years, microfluidics has demonstrated great potential for revolutionizing microbiome research (6). The most widespread applications include isolating and sorting microorganisms using droplet technologies or microwell technology and simulation of physical and chemical conditions associated with microorganism growth (7). Moreover, microfluidic is also used for proteomic and multiomic sample preparation. Diverse microdevices present controlled lysis capability using different modalities, including mechanical lysis, laser lysis, thermal lysis, and electrical lysis8. Some multilayer microfluidic devices, including the SciProChip, have been developed, allowing cell lysis and protein preparation for non-targeted protein identification (8). Efficient multiomics-based methods allowing nucleic acids and protein characterization have been recently developed. However, they are currently limited to mammalian cell-related research (5, 9, 10). These methods can be used to study biopsies, tissues or single cells. However, microorganisms are often studied at the population level rather than cellular level for reasons that include the limitations in mass spectrometry sensitivity and analysis time required per sample. Currently, the instrument time required for analyzing a single sample is ∼1 h per sample in most institutes having MS/MS acquisition rates of ∼10−20 Hz (11). Even if the new generation of mass spectrometers or labeling strategies reduce this time, it is more realistic to analyze field samples at the population level than at the single cell level since it is well adapted to provide an overview of the taxonomic and metabolic diversity of the samples. In our case, microfluidic devices are used because they can improve sensitivity and increase proteome coverage, for low abundant samples (12-14).

In a previous report (15), we proposed a ChipFilter microfluidic device as a miniaturized reactor for protein extraction and proteolysis directly from mammal cells. The device has two reaction chambers of 0.6 μl volume separated by an ultrafiltration membrane made using regenerated cellulose to concentrate or retain large polypeptides while releasing small molecules of less than 10 kilodaltons (kDa). The original protocol was modified to study multitaxonomic samples of microbial cells directly into the device to perform all steps necessary for sample preparation, from microbial cell lysis to proteolysis (14). To study proteins, we have shown that both for eukaryotic cells and microorganisms, sample preparation in the ChipFilter makes it possible to identify more proteins and more peptides than their preparation using conventional protocols (including FASP, in-gel, and in-solution proteolysis) for low-abundant samples. In addition to these improved analytical performances, the ChipFilter can be used for several analytical steps, including sample collection, handling of low sample density, automation, and effective cell lysis of a wide range of bacteria and fungi.

In the present work, we propose a new workflow to obtain a new benefit of the ChipFilter: the possibility to analyze proteins and nucleic acids simultaneously from the same sample. We performed lysis and proteolysis of microbial cells directly in the device. After peptide recovery and analysis, DNA from the same samples was recovered and successfully analyzed by PCR and sequencing. The workflow was applied to a complex mixture of known compositions as a proof of concept. Bottom-up proteomics led to the identification of more than 5000 proteins. DNA from the same sample was recovered and sequenced with a performance like DNA extracted with a commercial kit. Our results show that the ChipFilter can be used for sample preparation for proteomic and genomic analyses.

## 2. Results

### 2.1 Proteomic analyses are effective

Our previous report showed that it is possible to perform cell lysis and proteolysis in the ChipFilter microfluidic device, which allowed the identification of more proteins and peptides than conventional protocols (14). To confirm these results, a commercial mixture of 17 species (Zymo Research, ref D6331), including bacteria (Gram-negative and Gram-positive), fungi, and archaea, was analyzed after ChipFilter cell lysis and proteolysis. This gut standard mixture has a non-uniform cell number composition and contains species commonly found in the intestinal microbiota of humans. Peptides obtained after proteolysis of the three replicates of the commercial gut standard were analyzed by mass spectrometry.

These samples were treated as if the sample composition was unknown. Using the whole TrEMBL database, 3,017 proteins, and 5,052 peptides were identified. Among these identifications, only 1,058 proteins and 2,129 peptides belong to the 17 species in the mixture (Table 1).

**Table 1.** Number of proteins, peptides, and Peptide-Spectrum Matches (PSM) identified from the 17 species composing the mixture and the contaminants (“target”) and from other species (“others”), according to the databank used. Database search were done with 3 modifications only

|  | Number of proteins<br>IDs (target/others) | Number of peptides<br>IDs (target/others) | Number of PSM<br>(target/others) |
| --- | --- | --- | --- |
| Database 1 | 2,930 / 0 | 9,522 / 0 | 23,694 / 0 |
| Database 2 | 2,207 / 1,177 | 5,016 / 2,481 | 11,996 / 6,866 |
| Database 3 | 2,183 / 1,288 | 4,880 / 2,613 | 11,496 / 6,993 |
| Database 4 | 1,058 / 3,017 | 2,129 / 5,052 | 4,839 / 17,180 |

### 2.2 DNA usable for analysis can be recovered after cell lysis and proteolysis in the ChipFilter

To study microorganisms, obtaining nucleic acids from the same samples after proteolysis would be advantageous. To verify the feasibility of this approach, we performed lysis and proteolysis on a ChipFilter using 1E6 cells of *E. coli, S. cerevisiae*, and *B. subtilis*. The flow was reversed (Fig. 1A) for nucleic acids to be analyzed by PCR. Efficient amplification was obtained for the three species belonging to different phyla. These amplifications were like DNA amplification of DNA extracted with a commercial purification kit (Fig. 1B).

**Fig. 1.**
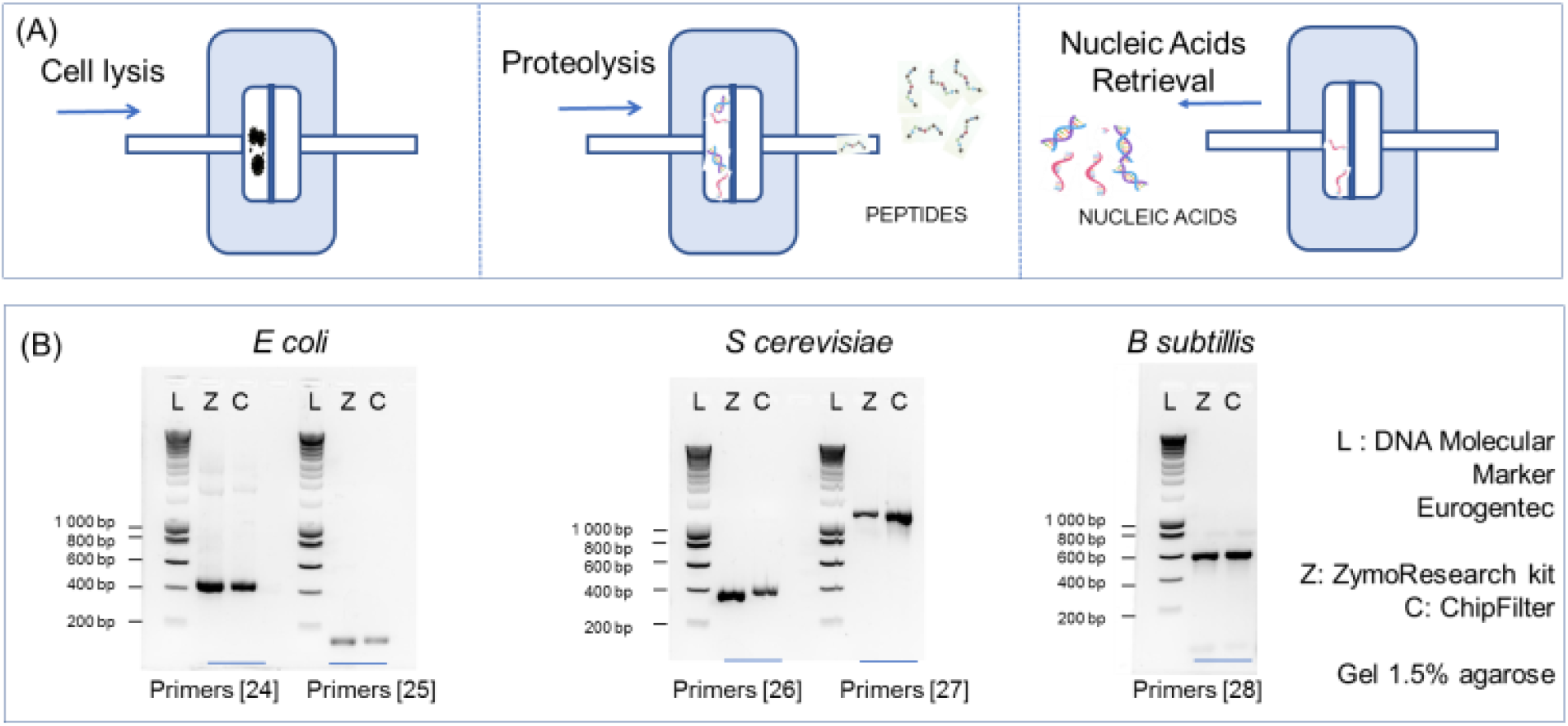
DNA can be recovered after ChipFilter proteolysis. (A) Experimental design. Cells are loaded and lysed in the ChipFilter (left). Peptides resulting from the proteolysis pass through the 10kD membrane to be analyzed by LC-MS/MS (middle). The flow is reversed, and the nucleic acids are recovered and can be used for DNA analysis (right). The blue arrow indicates the flow direction. (B) PCR analysis of DNA extracted from E coli (left), S cerevisiae (middle), and B subtilis (right) using the ChipFilter (C) or a commercial kit (Z). The sizes of the amplicons are equal to the expected sizes (16-20).

### 2.3 DNA can be efficiently sequenced

To further establish the multi-omics capability of the ChipFilter, the same commercial mixture of 17 species was sequenced after ChipFilter cell lysis and proteolysis. The benefit of using a commercial standard is that the sequencing results can be assessed qualitatively and quantitatively.

Sequencing experiments have made it possible to identify all the species. Even the genome of the less abundant species, *Clostridium perfringes*, estimated to be 400 cells/analysis according to the supplier’s data, could be sequenced after preparation in the ChipFilter (Fig. 2A). The relative abundances of the species were determined. These correlate well with the theoretical abundances given by the supplier, except for 3 species (*Saccharomyces cerevisiae, Candida albicans, Methanobacter smithii*) that were underestimated (Fig. 2A). The log-distributed abundances of the species are comparable to that obtained after DNA preparation using a commercial kit (Fig 2B), for which some species were also underestimated (*Saccharomyces cerevisiae, Candida albicans, Bifidobacterium adolescentis*).

**Fig. 2.**
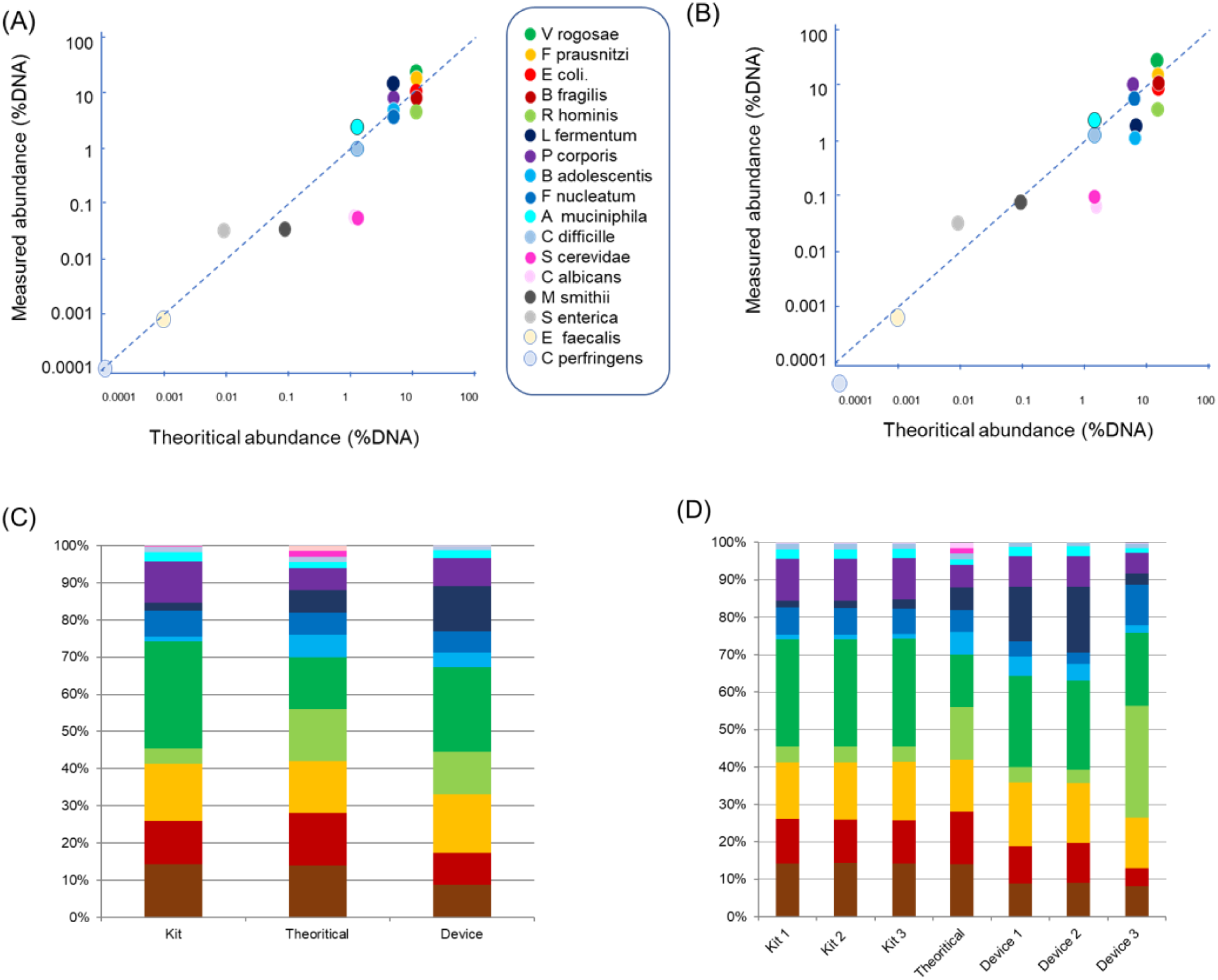
DNA from a defined mixture of microorganisms can be sequenced after ChipFilter lysis and proteolysis. (A) Log-distributed abundance (%DNA) after ChipFilter sample preparation vs theoretical abundance. (B) Log-distributed abundance (%DNA) after commercial kit sample preparation vs theoretical abundance (C) Average quantities of DNA from different species determined after extraction with a commercial kit (left) and extraction in the ChipFilter (right). (D) Quantities of DNA from different species were determined after extraction with a commercial kit (left) and extraction in the ChipFilter (right) for each replicate. The color code is the same for all the graphs

The relative abundance of the different species was determined for the three replicates treated either with the ChipFilter or a commercial kit. The difference with the theoretical distribution evaluated by the MIQ score is identical for samples extracted with the commercial kit (43.3 ± 0.6) and those processed on the ChipFilter (46.6 ± 4.6). Our results suggest that the ChipFilter allow to estimate the relative abundance of the species with a higher accuracy than the commercial kit (Fig 2C) but with a with lower precision (Fig. 2D), results being less reproducible among the replicates.

### 2.4 Use of Genomic data to improve proteomic data quality

As the DNA sequencing experiments were able to determine the sample taxonomic composition, we used this information to evaluate the effect of the database size on the quality of the proteomic identifications. This was done using four protein databases containing 4,098,824 to 249,752,136 sequences. Databank 1, contains only the protein sequences of the 17 species identified by DNA sequencing, exported from TrEMBL. The other three databases of increasing size are banks that can be used when there is no indication of the phylogenic composition of gut samples: Databank 2 comprises the sequences of abundant species of the gut microbiome, Databank 3 is a larger bank of the mouse gut microbiome species, and Databank 4 is the entire TrEMBL bank, version 02-2024. A contaminant database (trypsin, keratins, etc.) was added and considered in all database searches. The total number of peptide and protein identifications increases with the size of the database; however, the number of protein and peptide identifications from the species of the commercial mixture decreases in parallel (Table 1, Fig. 3).

**Fig. 3.**
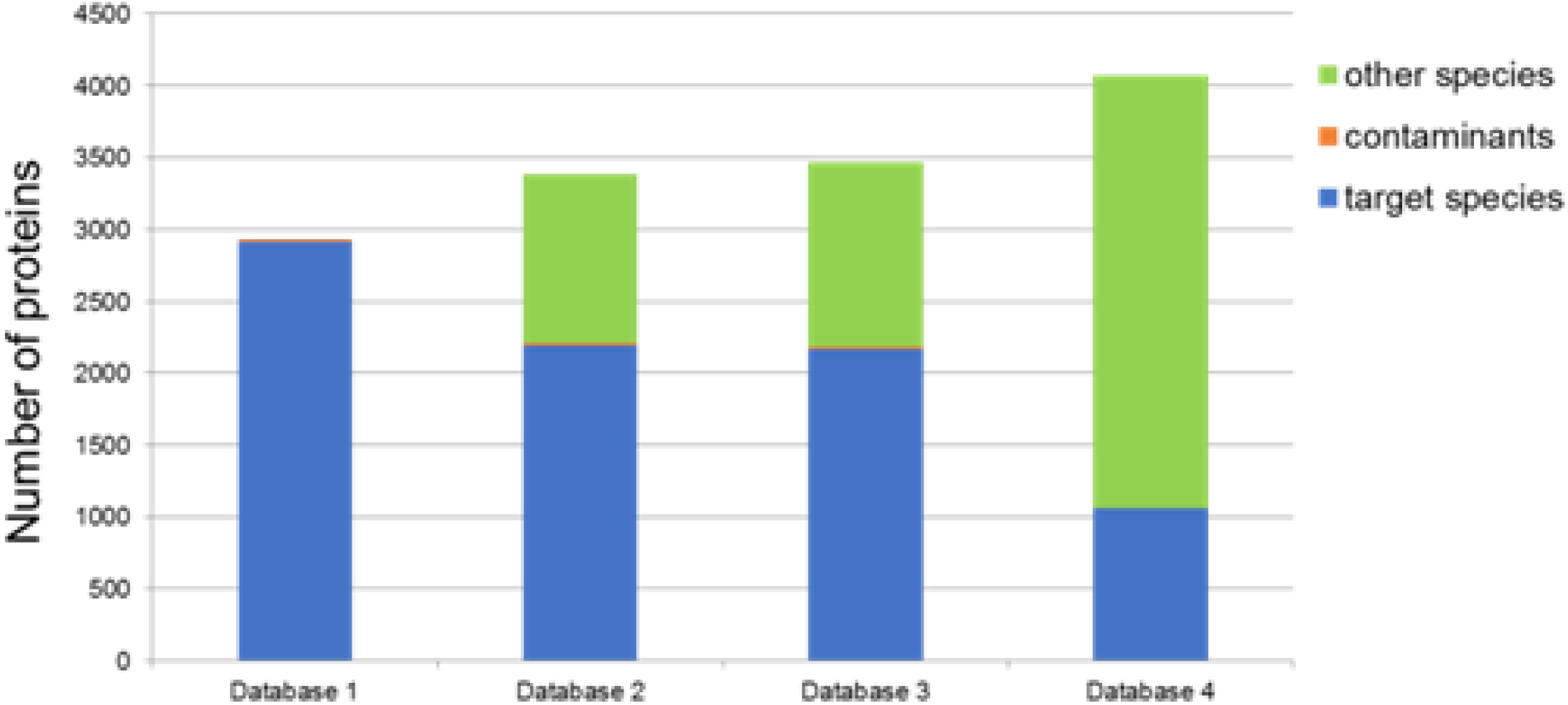
Several proteins were identified from the 17 species composing the mixture and from other species, according to the databank used.

The genomic analysis allows optimizing the proteomic database and improving post-translational modifications characterization. Using Database 1 (sequences of the 17 species, 6 modifications allowed), 5006 proteins and 16,137 peptides belonging to the 17 expected species were identified. The identification number per replicate was 2915 ± 412 proteins and 7046 ± 1272 peptides.

Some functional data were deducted from our analysis. The proteomic data were used to estimate the biomass of each species21. The results are compared with those obtained with theoretical DNA and cell number distribution (Figure 4A). The Gene Ontology could be determined for 3,891 identified proteins (Figure 4B). Some Post-Translational Modifications (PTMs) could be determined (Figure 4C), including 34,618 peptides with K acetylation, 98 with K succinylation, and 163 with Y phosphorylation.

**Fig. 4.**
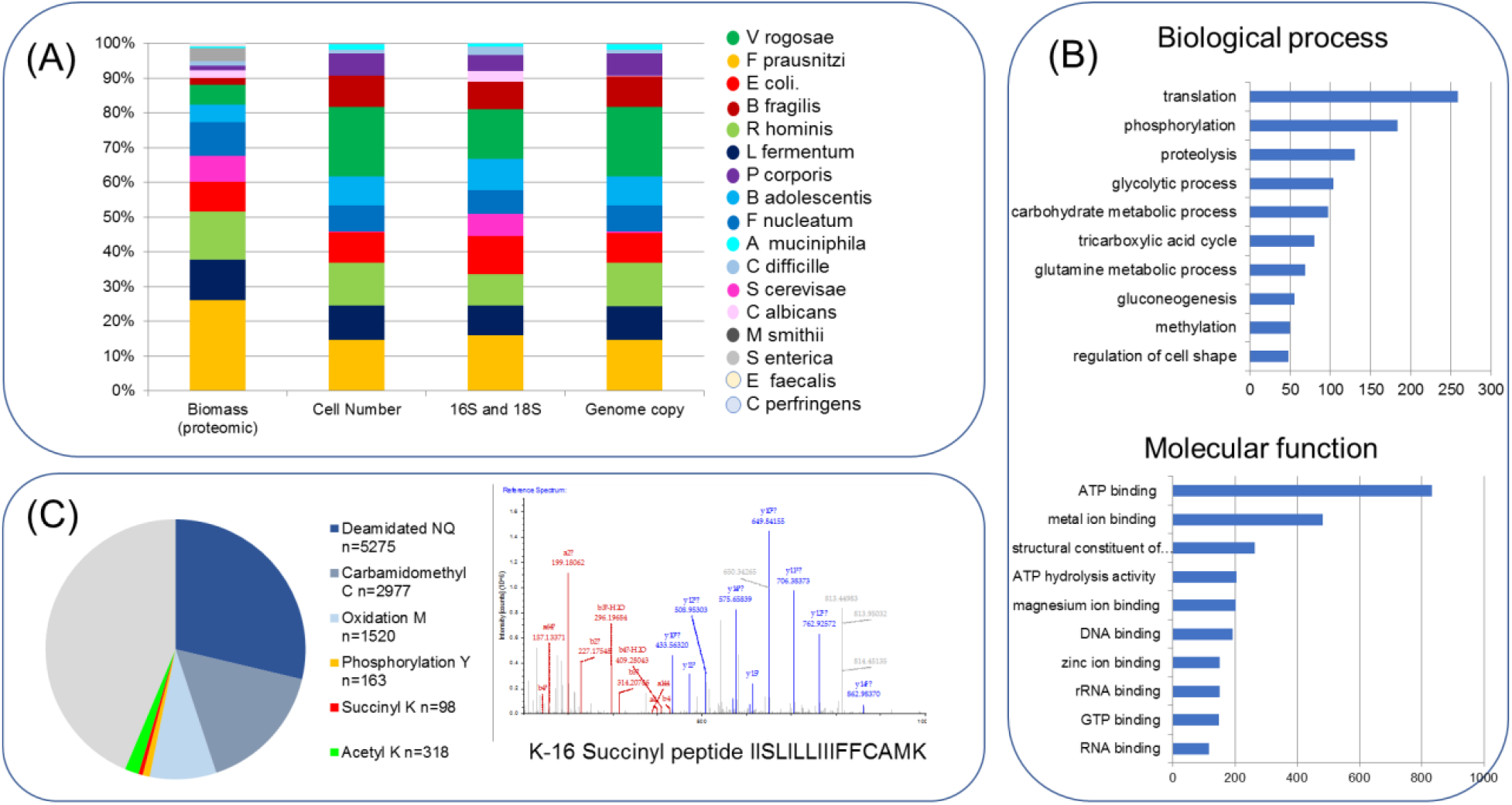
Examples of functional information provided by the current proteomic analyses. (A) Contribution to biomass (left) compared to the data provided by the manufacturer. (B) Gene Ontology annotation was determined for 3,891 proteins. (C). Identification of some PTMs. Left: distribution of modified peptides. Right: example of a modified peptide

## 3. Discussion

The present work aims to evaluate the possibility of studying DNA and proteins from the same samples after sample preparation in a microfluidic device. Microbial cells were lysed directly into the ChipFilter, and proteins were digested. Peptides were recovered and analyzed by nanoLC-MS/MS. Then, the flow was reversed, and the DNA was collected for further study. This workflow identifies more proteins than conventional protocols, as shown by previous work (14). The present study clearly shows that in addition it is also possible to amplify and sequence DNA, with results comparable to results obtained using the conventional extraction method.

Using PCR, we amplified the DNA of three model species after cell lysis and proteolysis in the ChipFilter. Commercial microorganism DNA preparation kits generally involve three steps: cell lysis, DNA purification, and removal of PCR inhibitors. Our positive PCR results confirm that the chemical cell lysis is effective (14). Moreover, it demonstrates that the regenerated cellulose membrane retains the DNA and that the filtration during the proteolysis step eliminates PCR inhibitors. Eventual material loss associated with non-specific absorption on the device or DNA degradation linked to the chemical reagents was not redhibitory to DNA analysis.

A more complex sample was then sequenced using the same extraction protocol. Our results show that the DNA of all the expected species in the mixture was sequenced, including the genome of the less abundant species, *Clostridium perfringes*, estimated to be 400 cells/analysis according to the supplier’s data. Relative abundances correlate well with the theoretical abundances, except for 3 species. The experimental distribution can be compared with the theoretical distribution using an index proposed by the supplier, the MIQ score. Considering this parameter, the DNA sequencing semi-quantitative results obtained with the ChipFilter are like those obtained with a commercial kit, even if the latter is more reproducible with the standardized commercial kit than those treated in the homemade ChipFilter. It can be attributed to our manual extemporaneous fabrication process for the ChipFilter device and the longer storage time of one of the replicates processed with the ChipFilter (supplementary fig S2).

We did not highlight any systematic identification bias linked to lysis in the ChipFilter. Misestimation of relative abundances determined by sequencing can have various causes. Some species, such as Saccharomyces cerevisiae and Candida albicans, are underestimated in our sequencing results. It is known that they are challenging to lyse without physical disruption 22,23 (Sup. Fig 1A). Another parameter is the cell number (Sup. Fig. 1B). In the mixture we used, both S. cerevisiae and C. albicans were taken at a low cell number. These two species also have a low GC content, which can impact the Illumina sequencing method (24) (supplementary figure 1C). It should be noted that this under-representation is also the case for the sample extracted with the commercial kit and is not specific to our protocol, suggesting that the ChipFilter extraction protocol does not induce any specific bias.

Proteomic analysis can provide functional information on the sample, complementing the genomic data. Since proteins convey structure and activities to cells, knowing their abundance provides a picture of cellular phenotypes on the molecular level. Proteomic studies can also provide information on the PTMs of proteins, some of which have been associated with the physio-pathological state of the microbiome. It is the case, for example, of bacterial protein acetylation that plays a prominent role in central and secondary metabolism, virulence, transcription, and translation (25), and has been associated with Crohn’s disease (26), succinylation which is associated with Quorum sensing system (27) or phosphorylation, involved in host-microbiome interaction (28), among many other possible effects. Because we carried out this work on a standard with no biological significance, as the microorganisms grew separately, we carried out these analyses only as a proof of concept, without biological interpretation. Even in our mixture, it was possible to annotate the function of 2,996 genes and to identify hundreds of PTMs, including acetylation, phosphorylation, and succinylation. Quantitative studies between several samples can also be carried out, including studies of enrichment in proteins involved in metabolic functions and enrichment in some PTMs. Other information can be obtained from proteomic analysis: differential analysis between samples, sample and protein clustering, protein interaction networks, or metabolic pathway analysis can be deduced from proteomic analysis (29).

The metaproteomic data generated to examine functions can also be used to analyze community structure. In most organisms, protein constitutes the largest amount of dry cellular material. Therefore, quantifying the total protein per species allows for assessing the biomass contributions of community members (21). Our results illustrate this possibility. Let us take the example of two species of very different cell number concentrations according to the manufacturer’s data, E. coli and S. cerevisiae, which represent 8.73% and 0.16%, respectively. Their respective contribution to the biomass estimated with proteomic data is 8.67% and 7.46%, respectively. This contribution is close to that obtained (8.73 and 6.42%) if the cell number contribution by their volume, estimated at 1 µm3 and 42 µm3 30-31, making the reasonable approximation that the biomass of a cell is proportional to the cell volume. This approximation is since most cells comprise about two-thirds water and that other components, like proteins, have a characteristic density of about 1.3 times the density of water (31).

Complementarity between approaches is also crucial from an analytical point of view. One of the difficulties with metaproteomic analysis is that the phylogenic composition of the sample is not known. Because proteomic identifications require sequence databases, it has been demonstrated that taxonomic and functional results strongly depend on the database choice (32). Increasing the database size to cover all possible sequences is attractive but potentially counterproductive for at least two reasons. First, the relationship between the database size and the number of Peptide-Spectrum Match (PSM) shows that the number of PSMs stops increasing at a certain point because the target PSM saturates when all peptides in the sample are identified. Decoy PSMs, for their part, are randomly identified because they happen to be hit by chance and thus increase in proportion to the database size. This effect affects the validation score threshold and leads to a decrease in peptide identification (33). We have not observed this phenomenon in our analysis. Second, many organisms have similar sequences in metaproteomics because of their phylogenic proximity. Increasing databank size increases the possibility that a PSM matches with a peptide from closely related species. That is why building a reference database through metagenomic sequencing is often a prerequisite for metaproteomics (34). In the present work, we compared the protein identifications obtained using a database derived from DNA sequencing with those obtained using three databases built with restrictive conceptions of the microbiome. The number of identifications increases with the size of the database, but the number of proteins attributed to the 17 species making up our standard decreases. These misinterpretations can lead to a bias in the functional interpretation of proteomic data. Culture contamination cannot be ruled out for some of the proteins identified from other species. For example, in the case of proteins from the Lachnospiraceae taxon, 213 proteins were identified using Database 2 (1929 PSM, representing 3.5% of the biomass). Another significant advantage of using a dedicated database is that it reduces search time and allows for more PTMs to be searched in a realistic time frame.

On the other hand, proteomic data can help with genome annotation. It has been estimated that only 52% to 79% of the average bacterial proteome coverage could be functionally annotated based on protein and domain-based homology searches, respectively. The experimental characterization is essential, particularly for accessory proteomes and understudied lineages (35).

Integration of omics data allows a better comprehension of a system. For example, diverse omics tools have been developed to guide the discovery and characterization of various microbial metabolites, which make it gradually possible to predict the overall metabolites for individual strains. The combinations of multi-omic analysis tools, such as OmicsNet (36) that combines genomic and proteomic data compensates for the shortcomings of current studies that focus only on single omics or a broad class of metabolites (37).

Therefore, the possibility of using a microfluidic system to carry out different analyses from a unique sample is particularly attractive. We plan to study further and adapt the ChipFilter protocol to analyze other molecules of interest, such as metabolites and RNA. Moreover, our strategy is easily up-scalable for larger samples or could, on the contrary, be down-scaled and adapted to be coupled with droplet microfluidics

## 4. Experimental

### 4.1 Materials

Octyl-β-D-glucopyranoside (ODG), protease inhibitor, dithiothreitol (DTT), ammonium bicarbonate (ABC), phosphate buffer saline (PBS), and iodoacetamide (IAM) were purchased from Sigma-Aldrich. Acetonitrile (ACN) and Formic Acid (FA) were purchased from Fisher Scientific. Trypsin Gold MS Grade was obtained from Promega and Lysozyme (50 ng/ml) from Thermo Fisher Scientific Eurogentec synthesized oligonucleotides. Qiagen provided Taq DNA polymerase and dNTP. DNA extraction kit (D4301) from Zymo Research was used.

Single microbial colonies of Saccharomyces cerevisiae, Escherichia coli, and Bacillus subtilis were used. Each of these model cells represents the fungi, Gram-negative, and Gram-positive bacterium types respectively. Luria-Bertani (LB) agar (Thermo Fisher Scientific) plates were inoculated with the cells and incubated at 37 °C overnight. Inoculum into LB broth was made and cells were cultivated until the optical density reached 1 at 600 nm. Before harvesting, cells were counted using a glass slide (Fisher Scientific) under a microscope. Collection of the cells was done by centrifugation at 2400 g for 5 minutes at room temperature. A single wash with phosphate-buffered saline (PBS) pH 7.4 was performed before pelleting and storage at -80 °C until further use.

A standard whole-cell mixture consisting of 21 representative strains from 17 gut microbiota species (Zymo Research, ref D6331) was divided into 10 aliquots and stored in the storage solution provided by the manufacturer at -80 °C. This standard contains 18 bacterial strains, including five strains of E. coli (JM109, B-3008, B-2207, B-766, and B-1109), 2 fungal strains, and 1 archaeal strain in staggered abundances, theoretically ranging from 20.01% to 0.0009% considering their cell number.

### 4.2 ChipFilter fabrication and use

The design and fabrication methodology of the microfluidic device has been explained previously (15). The ChipFilter is a PDMS device composed of two reaction chambers (inner diameter 4 mm, height 50 µm, volume 0,6 µL each) separated by a regenerated cellulose filtration membrane (diameter 4.3 mm) slightly larger than the reaction chamber. Twelve pillars of 150 µm diameter were designed inside the two chambers to maintain the filtration membrane.

A 3D master mold was used to ensure proper integration of the membrane. The patterns were designed with CleWin5 software and printed at high resolution (25,400 dpi) on a photosensitive film by a photoplotter FilmStar-PLUS:2 mask for each reaction chamber, with a 2-step lithography for 3D molds. The mold comprises 2 layers of negative photoresist on the silicon wafer. The first layer is a SU-2007 resin (8 µm thickness, spin-coated at 2000 rpm), dedicated to the structure to incorporate the filtration membrane and to avoid any leakage of the reaction chamber. After baking at 95°C for 2 minutes, the first mask features were transferred onto the wafer by photolithography using a UV-KUB3 aligner (LED, 40 mW/cm2, 160 mJ/cm2, 5 seconds). After insolation through mask 1 and post-baking at 95°C for 2 minutes, a second photoresist layer was spin-coated on the first. The second layer is a SU-2050 resin (50 µm thickness, spin-coated at 4080 rpm) dedicated to reaction chambers and pillars. Mask 2 is aligned with the patterns of mask 1 using a UV-KUB3 aligner. The wafer was then exposed to LED light (40 mW/cm2, 160 mJ/cm2, 4 s). Finally, the developer removed the non-exposed part of the photoresist and revealed the 3D patterns.

PDMS elastomeric polymer (Sylgard 184) has been used for precise replica molding. A two-component mixture (base/curing agent, 10:1 (w/w)) liquid pre-polymer was poured on the mold and cured at 70°C for 1 h. After exposure to an air plasma (20 W, 8 sccm O2 flow at 0.13 mbar) for 1 min, the 2 PDMS parts were assembled with the membrane between them after the membrane was positioned on the first chamber covering the first reaction chamber. The alignment of the two chambers was controlled under a microscope using alignment marks. A high-pressure contact between the 2 PDMS parts will allow a tight sealing around the filtration membrane. The entire assembly was placed at 90°C for at least 15 minutes to enhance bonding.

### 4.3 Sample preparation in the ChipFilter

Microbial lysis was realized directly on the ChipFilter without any pre-treatment as described14. Briefly, 30 μL of cells were introduced into the ChipFilter using a piston syringe (Agilent) and syringe pump (Harvard Apparatus), maintaining a flow rate of 0.01 ml/minute. For the sequential injection of lysis buffer [0.5mg/mL Lysozyme in Working Buffer (WB) 1% (w/v) ODG, protease inhibitor in 150 mM Tris-HCl pH = 8.8], 20 mM DTT in WB buffer, 50 mM IAM in WB and 50-mM ABC buffer was achieved using a flow-EZ pressure module, flow controller, M-switch (Fluigent), and the software Microfluidic Automation Tool (Fluigent). The flow rate and volume were maintained in two stages at 2 μL/min for 45 μl and 1 µL/min for 30 μL with the upper-pressure limit at 900 mbar. Finally, proteolysis was performed at room temperature by introducing 20 µL of trypsin (final concentration of 0.1 µg/µL in 50 mM ABC). A constant flow of 50 mM ABC was maintained for 150 min to ensure the mixing of the proteins with trypsin. The resulting proteolytic peptides contained in the flowthrough passing through the filter were directly transferred to the sample loop of the nanoLC system before being finally concentrated in a trapping column (C18 Pepmap, 300 μm i.d. × 5 mm length, ThermoFisher Scientific).

In the case of the standard gut microbiota analysis, three replicates of 75 μl of the mixture evaluated to 3.94E8 cells by the manufacturer were thawed in ice. Cell lysis was performed with lysis buffer supplemented with 0.5 mg/ml lysozyme. All the subsequent steps were done as described above.

After proteolysis, the flow was reversed, and the nucleic acids were recovered in 35 µL water.

### 4.4 Protein analysis

Peptides Liquid Chromatography tandem Mass Spectrometry (nanoLC-MS/MS) analysis. Samples were analyzed by nanoLC-MS/MS using an RSLC nano U3000™ system coupled to a nanoESI Q-Exactive HF mass spectrometer (ThermoFisher Scientific).

Peptides were desalted and transferred from the trap column to be separated on a capillary reverse-phase column (Acclaim™ PepMap™ RSLC C18, 2 μm, 100Å 100 Å, 75 μm i.d. × 50 cm, ThermoFisher Scientific) at 45°C with a linear 120 min gradient elution from 2.5% to 60% of solvent B (water/ACN 10:90 (v/v), 0.1% FA) in solvent A (water/ACN 98:2 (v/v), 0.1% FA) at a 220 nL/min.

Samples were analyzed in Top20 data-dependent acquisition (DDA) high-energy dissociation (HCD) mode. One nanoESI (1.6 kV) full MS survey (m/z range of 375-1500, resolution of 60,000, automatic gain control (AGC) 3E6, maximum injection time 60 ms) was followed by up to 20 MS/MS acquisition on the most 2+ to 5+ charged ions (normalized collision energy 28, precursor isolation window 2 m/z, resolution 15,000, AGC 1E5, max injection time 60 ms, minimum MS2 target value 1E3 and dynamic exclusion for 20 s).

Proteomic Data Analysis. Spectra were processed using Proteome Discoverer v2.4 (ThermoFisher Scientific) interfaced with the Mascot search engine. Four sequence databases of various sizes were used (Table 2), complemented with a databank of current laboratory contaminants (trypsin, keratins, and others).

**Table 2.**
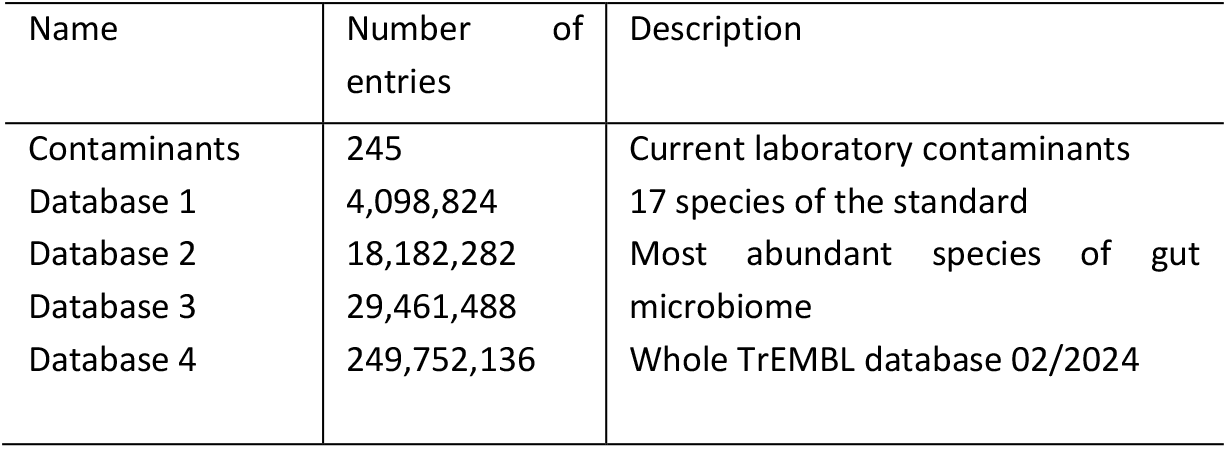
Protein databases are used for protein identification.

The database search was performed with the following parameters: MS and MS/MS mass tolerance 10 ppm and 0.02 Da, respectively, and trypsin specificity with up to 2 missed cleavages. The following partial amino acid modifications were considered: carbamidomethylation (C), deamidation (NQ), and oxidation (M). For databank 1 (containing only the sequences of the 17 species of the mix), an additional search including succinyl and acetyl (K) and Phospho (Y) together with the 3 others PTM was also performed. Proteins with at least one high-confidence peptide with more than 6 residues were validated. Target FDR was set at 0.01.

The same parameters were used to assess the effect of databank size. Due to the size of some banks, only three partial amino acid modifications were allowed: carbamidomethylation (C), deamidation (NQ), and oxidation (M), and searches were done directly on Mascot, without using Proteome Discoverer again to avoid any bias in the raw data processing.

### 4.5 Nucleic acid analysis

PCR analysis. DNA from three microbial species, B subtilis, E coli, and S cerevisiae, was used for PCR amplification. The DNA was extracted after ChipFilter preparation or using a commercial kit (ZymoBIOMICS DNA microprep kit, ref D4301, ZymoResearch) according to the manufacturer’s recommendations. The workflow of this kit, which is designed to extract DNA for microbiome analyses, includes microorganism lysis by bead beating, DNA purification using a spin column, and PCR inhibitor elimination using filtration. Species-specific primer sequences were designed according to the literature (Table 3) (24-28). The optimal PCR conditions were determined experimentally and were as follows: 3 min denaturation at 94° followed by 30 cycles of 94°C for 30 sec, annealing temperature (Table 3) for 30 sec and 30 sec at 72°C and 10 min at 72°C. The thermal cycler was from Analytic Jena. The amount of crude DNA extract was 2.5 µL for 50 µL of PCR mixture. Three µL of the amplified product was analyzed by electrophoresis on 1.5% agarose gels.

**Table 3.**
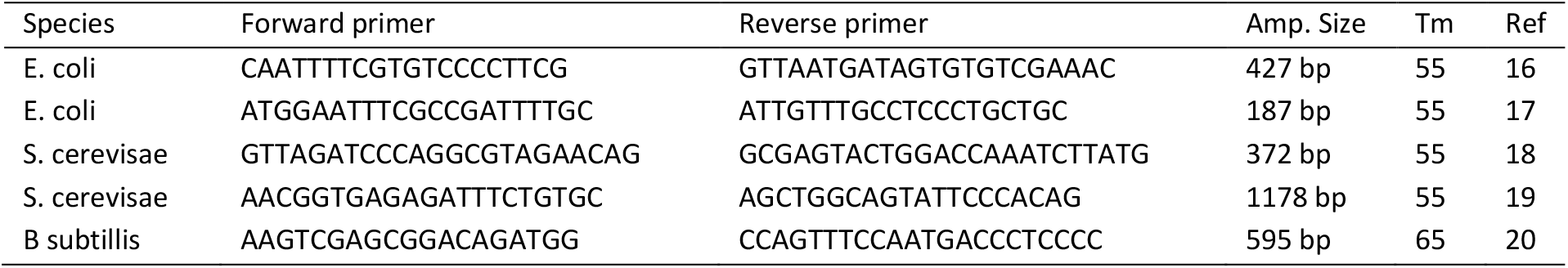
Primer sequence and annealing temperature used for the PCR experiments. Tm: annealing temperature

DNA sequencing. DNA extracted from the standard gut microbiota with a commercial kit (3 replicates) and with the ChipFilter (3 replicates) was sent to the Plateforme de Genotypage/Séquençage of the Paris Brain Institute (Paris, France) for sequencing. Library and multiplexing preparation were done using the Illumina DNA Prep kit (Illumina), which uses a transposon system. The quality of the library was assessed using Tapestation and fluorometric quantification. The library was sequenced using a Nextseq 2000 Illumina sequencing apparatus.

Barcode trimming was realized using Trimmomatic. Forward and Reverse reads were aligned and mapped on the reference genome using Bowtie238. The coverage for each genome was then determined (https://github.com/mmarbout/chipfilter).

Quality control of the sequencing quality was assessed using an MIQ score proposed by Zymo Research (https://github.com/Zymo-Research/miqScoreGutStandardShotgun)

## 5. Conclusions

The ChipFilter is a versatile device that allows protein and DNA analysis from the same sample aliquot. This system is particularly well suited to small-volume samples, for which complementary information can be obtained. In the future, we believe this versatile tool could be applied to additional omics studies to characterize a single sample aliquot even deeper.

## Author Contributions

RK RK, MN, and YV designed and performed the experiments. RK RK, JV, and YV analyzed the proteomic results. IH, MM, and YV analyzed the genomic results. JV supervised the work and funding. YV and RK wrote the original draft of the paper; all the authors had a critical review of the published work.

## Conflicts of interest

There are no conflicts to declare. Data availability

The mass spectrometry data have been deposited on the ProteomeXchange Consortium via the PRIDE partner repository with the dataset identifier PXD051944.

The github for DNA sequencing result analysis is available at: https://github.com/mmarbout/chipfilter

## Acknowledgments

This project has received funding from the European Union’s Horizon 2020 research and innovation program under the Marie Skłodowska-Curie grant agreement no. 754387. NanoLC-MS/MS equipment was subsidized by Conseil Régional d’Île-de-France (Sesame N°10022268).

This work has benefited from the technical contribution of the joint service unit CNRS UAR 3750 for microfluidic device fabrication. Sequencing was realized at the Plateforme de Genotypage/Séquençage of the Paris Brain Institute (CNRS UMR 7225 – Inserm U 1127 – Sorbonne Université UM75). The authors would like to thank the personnel of these units for their kind advice and support during the development of the experiments.

## Notes

https://github.com/mmarbout/chipfilter

### Competing Interest Statement

The authors have declared no competing interest.

