## Supplementary for "A single microfluidic device for multi-omics analysis sample preparation"

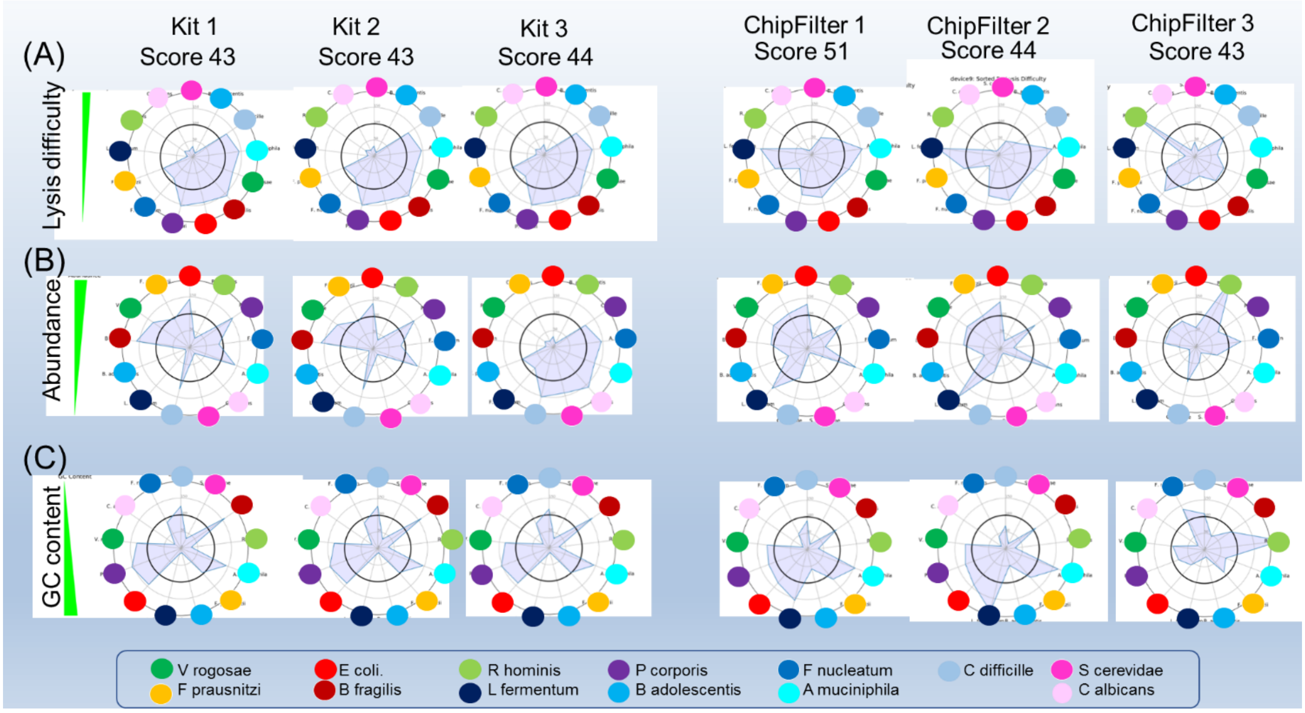


Supplemental figure S1. Deviation from the expected relative abundance for the samples extracted with the commercial kit (left) and the ChipFilter (right). On the radars, species have been ranked according to the difficulty of the lysis (A), the abundance (B), and the GC content (C).

### A. Device fabrication

**Internal structure**

1, 2, and 3 indicate pillar structures, alignment marks, and filtration membrane

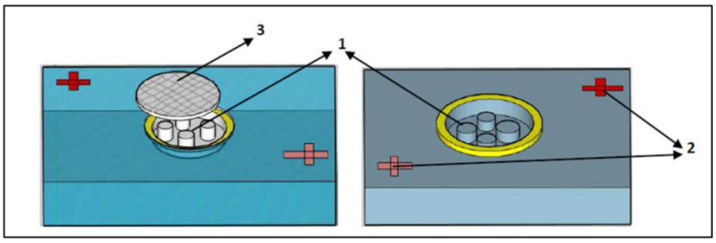


**External structure**

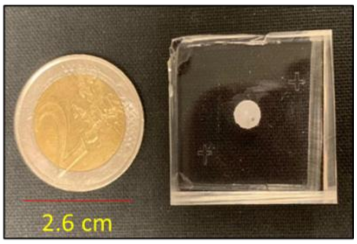


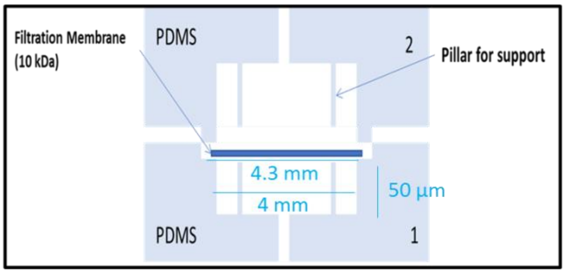


**Dimensions**
Reaction chamber diameter – 4mm
Membrane diameter – 4.3mm
Length of the chamber - 50µm

**(Ndiaye et al., 2020)**

The design and fabrication methodology of the microfluidic device has been explained previously7. The ChipFilter is a PDMS device composed of two reaction-chambers (inner diameter 4 mm, height 50 µm, volume 0,6 µL each) separated by a regenerated cellulose filtration membrane (diameter 4.3 mm) slightly larger than the reaction chamber. Twelve pillars of 150 µm diameter are designed inside the two chambers to maintain the filtration membrane.

A 3D master mold is used to ensure proper integration of the membrane. The patterns were designed with CleWin5 software, printed at high resolution (25,400 dpi) on a photosensitive film by a photoplotter FilmStar-PLUS: 2 masks for each reaction chamber, with a 2 steps lithography for 3D molds. The mold is composed of 2 layers of negative photoresist on the silicon wafer. The first layer is a SU-2007 resin (8 µm thickness, spin-coated at 2000 rpm), dedicated to the structure to incorporate the filtration membrane and to avoid any leakage of the reaction chamber. After baking at 95°C for 2 minutes, the first mask features were transferred onto the wafer by photolithography using a UV-KUB3 aligner (LED, 40 mW/cm2 , 160 mJ/cm2, 5 seconds). After insolation through the mask 1 and post baked at 95°C for 2 minutes, a second photoresist is spin-coated on the first one. The second layer is a SU-2050 resin (50 µm thickness, spin-coated at 4080 rpm) dedicated to reaction chambers and pillars. The mask 2 is aligned to the patterns of the mask 1 using a UV-KUB3 aligner. The wafer is then exposed at LED light (40 mW/cm2, 160 mJ/cm2, 4 s). Finally, the developer removed the non-exposed part of the photoresist and reveal the 3D patterns. PDMS elastomeric polymer has been used to replicate the features from the mold with high precision.

PDMS Sylgard 184 is used for the replica molding. Using a two-component mix (base/curing agent, 10:1 (w/w)), the liquid pre-polymer is poured on the mold and cured at 70°C for 1 h. After being exposed to an air plasma (20 W, 8 sccm O2 flow and 0.13 mbar pressure) for 1 min, the 2 PDMS parts are put in contact with the membrane between them. The membrane is deposited on the first chamber before the 2 PDMS parts assembly. It covers the first reaction chamber and the alignment of the two chambers is obtained under a microscope using alignment marks. A high-pressure contact between the 2 PDMS parts will allow a tight sealing around the filtration membrane. The whole setup is placed at 90°C for at least 15 minutes to enhance bonding.

### B. Experimental setup

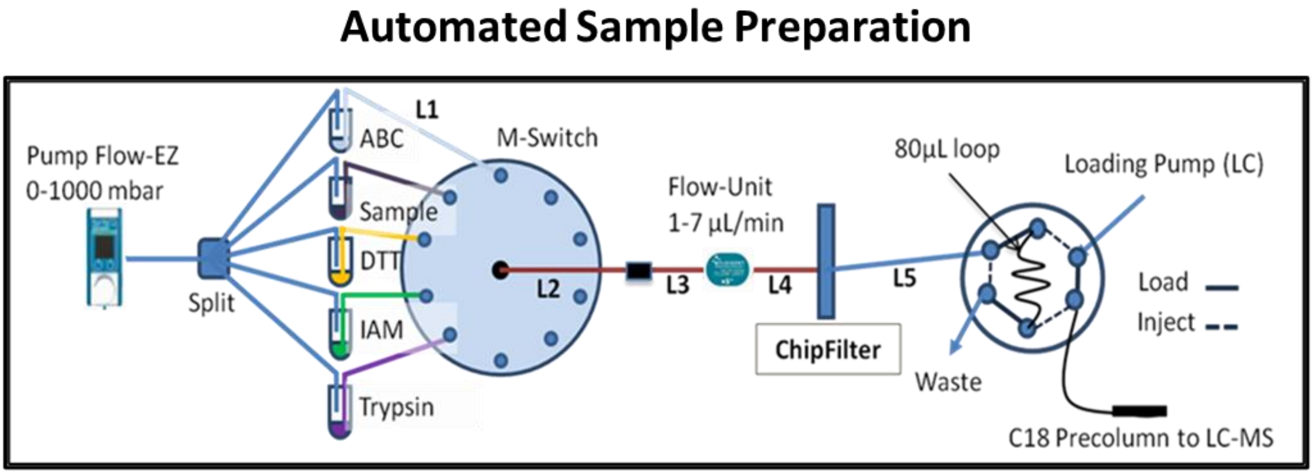


*Ndiaye et al., 2020*

Samples were introduced into the ChipFilter using a piston syringe (Agilent) and syringe pump (Harvard Apparatus) maintaining a flow rate of 0.01 ml/minute.

For the sequential injection of lysis buffer [0.5mg/mL Lysozyme in Working Buffer (WB) 1% (w/v) ODG, protease inhibitor in 150 mM Tris-HCl pH = 8.8], 20 mM DTT in WB buffer, 50 mM IAM in WB and 50-mM ABC buffer was achieved using a flow-EZ pressure module, flow controller, M-switch (Fluigent), and the software Microfluidic Automation Tool (Fluigent). The flow rate and volume were maintained in two stages at 2 µL/min for 45 µl and 1 µL/min for 30 µL with the upper-pressure limit at 900 mbar. Finally, proteolysis was performed by introducing 20 µL of trypsin (final concentration of 0.1 µg/µL in 50 mM ABC) at room temperature. A constant flow of 50 mM ABC was maintained for 150 minutes, to ensure the mixing of the proteins with trypsin. The resulting proteolytic peptides contained in the flowthrough passing through the filter were directly transferred to the sample loop of the nanoLC system before being finally concentrated in a trapping column (C18 Pepmap, 300 µm i.d. × 5 mm length, ThermoFisher Scientific).

In the case of the standard gut microbiota analysis, three replicates of 75 µl of the mixture evaluated to 3.94E8 cells by the manufacturer were thawed in ice. Cell lysis was performed with lysis buffer supplemented with 0.5 mg/ml lysozyme. All the subsequent steps were done as described above.

After cell lysis and proteolysis, the flow was reversed, and the nucleic acids were recovered in 35 µL water.

### C. Experimental design

| | |
|---|---|
| Sample | ZymoReasearch gut standard<br>Aliquots of 75 µL (≈3.94E8 cells) stored at -80°C |
| ChipFilter | |
| Replicate 1 | Lysis, Protein and DNA extraction : D1<br>Protein analysis : D2<br>DNA analysis : D105 (DNA strored at -80 until use) |
| Replicate 2 | Lysis, Protein and DNA extraction : D2<br>Protein analysis : D3<br>DNA analysis : D105 (DNA strored at -80 until use) |
| Replicate 3 | Lysis, Protein and DNA extraction : D90<br>Protein analysis : D91<br>DNA analysis : D105 (DNA strored at -80 until use) |
| Commercial kit | |
| Replicate 1 | Lysis, DNA extraction: D30<br>DNA analysis : D105 (DNA strored at -80 until use) |
| Replicate 2 | Lysis, DNA extraction: D30<br>DNA analysis : D105 (DNA strored at -80 until use) |
| Replicate 3 | Lysis, DNA extraction: D30<br>DNA analysis : D105 (DNA strored at -80 until use) |

### D. Analytical performances

| | | |
|---|---|---|
| Device characteristics | | |
| External Size | 3 cm x 3 cm x 1cm | Ndiaye et al., 2020 |
| Reaction Chamber | 4mm diameter x 50 µm | |
| Filtration membrane | 10 kDa nitrocellulose | |
| Working volume | 10-200 µL injected | |
| Operating modes | On-line (coupling with precolumn) and off-line (vials) | |
| Protein analysis | | |
| Analysis time (lysis, proteolysis) | 377 min | Ravi Kumar et al., 2024 |
| Sensitivity E coli 10E2 cells | 163 ±18 proteins ; 1 162 ±126 peptides | |
| Sensitivity E coli 10E6 cells | 1,999 ±54 proteins ; 9 770 ±108 peptides | |
| DNA analysis | | |
| Quantity[a] | 39.4 ±22.4 ng/µL in 30 µL | Present paper |
| Possible uses | PCR, Illumina Sequencing | |

A: determined using Nanodrop after lysis and proteolysis of 75µL of gut standard microbiome (ZymoResearch, ref D6331)

### E. Quality controls for proteomic analyses

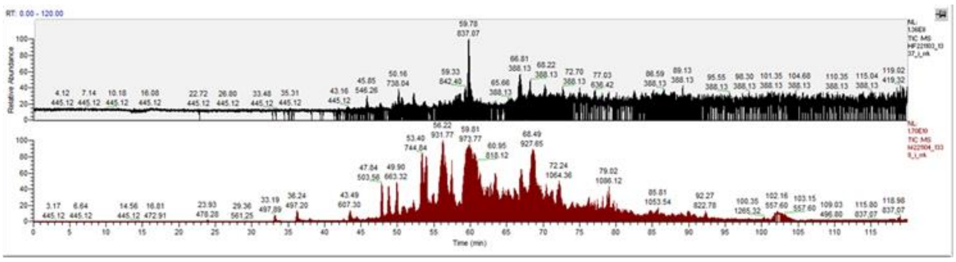

Blank 1
TIC 1.36E8

Replicate1
TIC 1.70E10

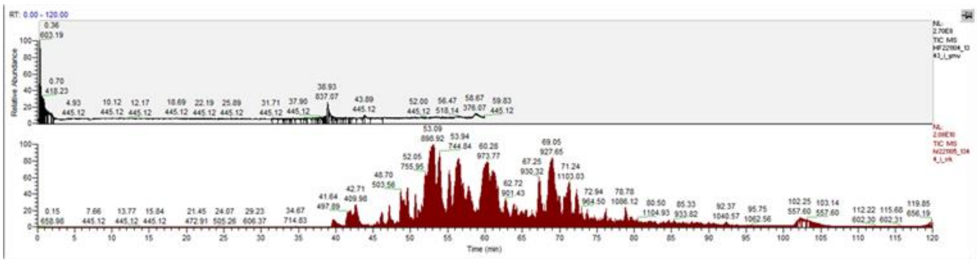

Blank 2
TIC 2.70E8

Replicate2
TIC 2.08E10

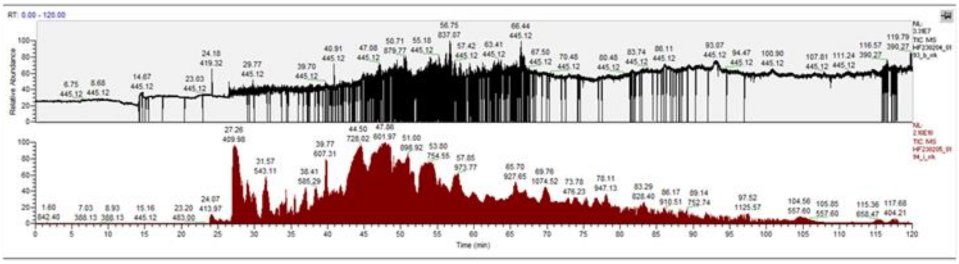

Blank 13
TIC 3.37E7

Replicate3
TIC 2.10E10

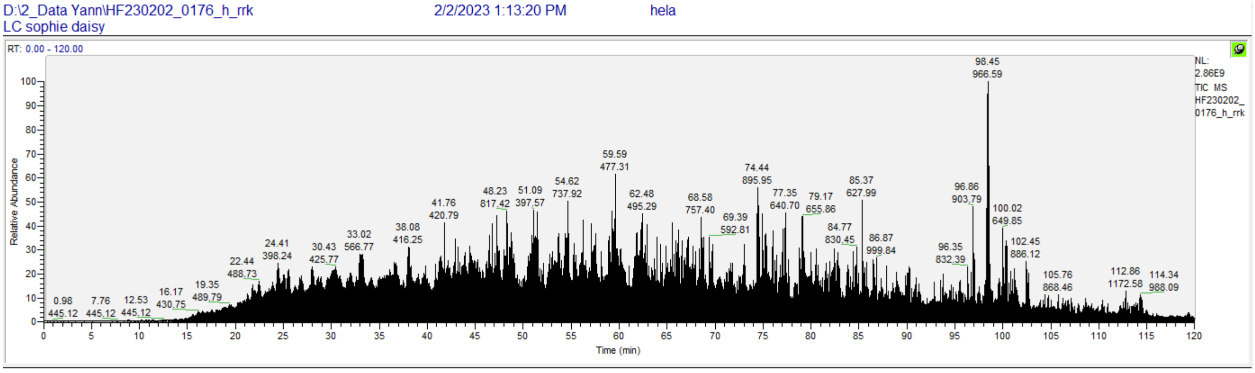


Due to the difficulty of cleaning the filter-based PDMS device and their low manufacturing cost, they are single-use devices.

For the mass spectrometer analysis, two controls are used:

-positive control: a mixture of 20 ng of peptides from hela cells are used. Analyzes were validated only if these controls are validated in terms of chromatographic profile and identification number reach at least 3000 proteins.

-negative control: to assess possible carry-overs between the samples, blanks are run between the samples

### F. Quality control of genomic analyses

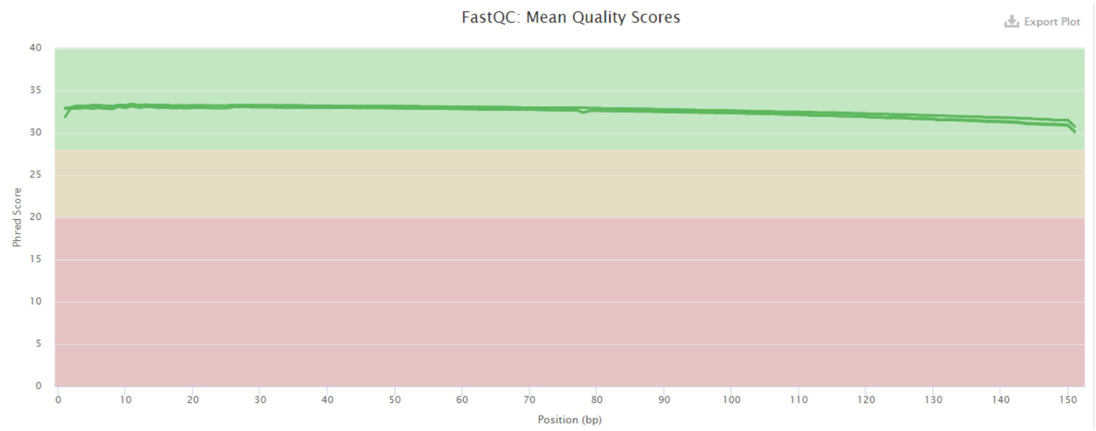


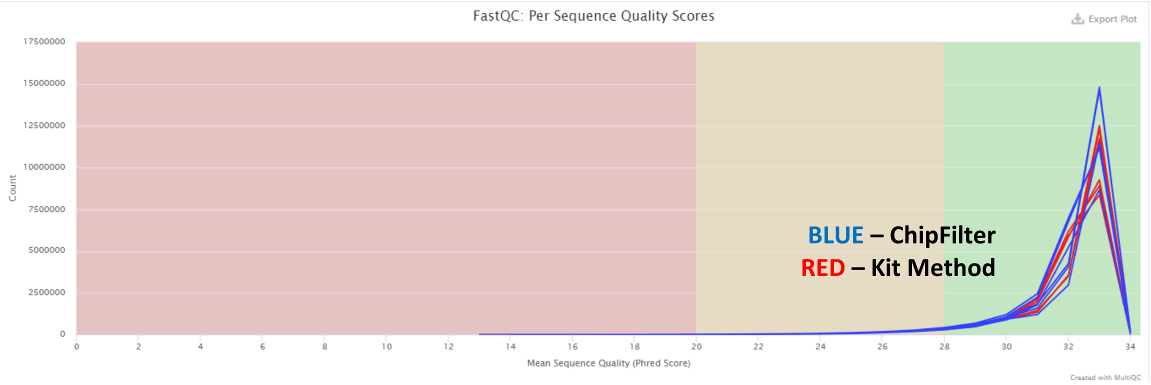


| Sample Name | % GC | Read Length | M Seqs |
|---|---|---|---|
| Kit1 Forward | 45% | 151 bp | 19.5 |
| Kit1 Reverse | 46% | 151 bp | 19.5 |
| Kit2 Forward | 45% | 151 bp | 19.8 |
| Kit2 Reverse | 45% | 151 bp | 19.8 |
| Kit3 Forward | 46% | 151 bp | 18.9 |
| Kit3 Reverse | 46% | 151 bp | 18.9 |
| Device1 Forward | 48% | 151 bp | 18.0 |
| Device1 Reverse | 48% | 151 bp | 18.0 |
| Device2 Forward | 48% | 151 bp | 23.7 |
| Device2 Reverse | 48% | 151 bp | 23.7 |
| Device3 Forward | 45% | 151 bp | 22.9 |
| Device3 Reverse | 45% | 151 bp | 22.9 |

Quality control of DNA sequencing analysis was realized using the MultiQC v1.12 software.

No difference were observed on the FastQC Mean and Per Sequence Quality Score between the sample extracted with the commercial kit and the sample extracted in the ChipFilter.
